# MEIOB and SPATA22 resemble RPA subunits and interact with the RPA complex to promote meiotic recombination

**DOI:** 10.1101/358242

**Authors:** Jonathan Ribeiro, Pauline Dupaigne, Clotilde Duquenne, Xavier Veaute, Cynthia Petrillo, Carole Saintomé, Orestis Faklaris, Didier Busso, Raphaël Guerois, Scott Keeney, Devanshi Jain, Emmanuelle Martini, Gabriel Livera

## Abstract

Homologous recombination is a conserved DNA repair process mandatory for chromosome segregation during meiosis. RPA, a ubiquitous complex essential to recombination, is thought to play a similar role during mitotic and meiotic recombination. MEIOB, a meiosis-specific factor with unknown molecular function, ressembles a RPA subunit. Here we use *in vivo* approaches to show that in mouse spermatocytes, DMC1 and RAD51 appear to be normally loaded in the absence of MEIOB but are prematurely lost from unrepaired recombination sites. This loss correlates with an accumulation of the BLM helicase on meiotic chromosomes. We also show that MEIOB alters the immunodetection of RPA subunits at meiotic recombination sites. Using electron microscopy and purified proteins, we demonstrate that the MEIOB-SPATA22 complex associates with and modifies the conformation of RPA-coated ssDNA. Finally, we identify structural homology between MEIOB, SPATA22 and RPA subunits, and show that MEIOB and SPATA22 interact through C-terminal OB-fold containing domains (OBCDs) like RPA subunits. Moreover, MEIOB and SPATA22 cooperate to interact with RPA through their OBCDs. Our results suggest that MEIOB, SPATA22 and RPA work together to ensure proper processing of meiotic recombination intermediates.

## Introduction

Meiosis is a specialized cell division program that halves the diploid genome of germ cells to produce haploid gametes. In most organisms, this requires the formation of a physical link between homologous chromosomes to ensure their segregation during the first meiotic division (1). These physical links are formed by reciprocal exchange of chromosomal arms, or crossovers (COs), which result from repair of DNA double-strand breaks (DSBs) by homologous recombination (HR) (2). While meiotic and mitotic HR share common features and factors they differ in several aspects (3). Meiotic HR is initiated by programmed DSBs (4, 5) and favors the use of homologous chromosomes rather than sister chromatids as repair templates (6). Only a subset of DSBs form COs; remaining DSBs are repaired without reciprocal exchanges and are important for pairing homologous chromosomes (7). Meiotic recombination is tightly linked to formation of the synaptonemal complex (SC), a tripartite structure comprising two lateral and one central (or axial) elements that stabilize the pairing of the homologs. These events are established during meiotic prophase I which is subdivided into four stages (leptotene, zygotene, pachytene, diplotene)(1). The specific requirements of meiotic recombination are satisfied by the combined action of meiosis-specific and ubiquitous proteins (6).

Single-stranded DNA (ssDNA) is generated at several steps during HR (8). After DSB formation, ends are resected to form 3’-ssDNA tails, which are then coated by the RecA-family recombinases to promote invasion of homologous sequences. This invasion induces displacement of the noncomplementary strand of template DNA, thus forming a displacement loop (D-loop) that can either be reversed or stabilized. These dynamic processes are under the control of numerous factors, including ssDNA-binding proteins and DNA helicases (9). During mitotic HR, the replication protein A (RPA) complex binds resected DNA to prevent degradation and secondary structure formation (10–12). Next, RPA is replaced by the RAD51 recombinase to form the presynaptic recombinogenic filament (13). During strand invasion, RPA can also stimulate RAD51-dependent strand exchange by coating and stabilizing the D-loop to prevent its reannealing (14, 15).

RPA is thought to play similar roles during mitotic and meiotic HR. Although RPA has been observed on unsynapsed chromosomes in mammals, only few faint RPA foci are detected when numerous, bright RAD51 foci have already accumulated on chromosome axes (16–18). Therefore, it is unclear whether RPA is loaded before or during the formation of the meiotic recombinogenic filament. In addition, the presence of numerous RPA foci during zygotene/pachytene stages of meiotic prophase, when a large fraction of DSBs have progressed through repair, suggests that RPA also plays a role in D-loop stability during meiotic HR (18, 19). In most organisms, in addition to RAD51, meiotic HR also requires the meiosis-specific recombinase DMC1 (20–22). The current model based on studies performed in *Saccharomyces cerevisiae* and *Arabidopsis thaliana* proposes that during meiosis, RAD51 and DMC1 can be loaded together onto broken ends and that RAD51 plays an accessory role in promoting DMC1 recombinase activity (23–25). This model has not been tested in mice, and the nature and dynamics of the recombinogenic filament during mammalian meiosis remain poorly understood.

RPA is an essential, evolutionarily conserved trimeric complex composed of RPA1, RPA2 and RPA3; it is involved in DNA replication, repair and HR. Recently, we and another group identified MEIOB (MEIosis specific with OB domains), a conserved meiotic paralog of RPA1 (19, 26). In mice, the absence of MEIOB induces an arrest at a zygotene/pachytene-like stage with an accumulation of unrepaired DSBs (19, 26). Several lines of evidence suggest that during meiosis, MEIOB forms a complex with SPATA22, a meiosis-specific factor with unknown function (26–29). Moreover, MEIOB and SPATA22 have been shown to interact with and colocalize with RPA subunits; the role and mode of these interactions remain to be understood (19, 26, 29).

Here, we demonstrate that MEIOB and SPATA22 share structural features and interaction patterns with RPA. We show that MEIOB and SPATA22 together condense ssDNA that is precoated with RPA *in vitro*, and alter immunodetection of RPA subunits *in vivo*. We reveal that in the absence of MEIOB, the BLM helicase accumulates on RPA-associated HR intermediates and that this accumulation is concomitant with DMC1 recombinase dissociation. These results suggest that MEIOB is essential for the proper processing of ssDNA-containing HR intermediates.

Altogether our data provide new insights into the properties and roles of the essential meiosis-specific proteins MEIOB and SPATA22, and our findings suggest that the MEIOB-SPATA22 complex modifies properties of RPA-coated ssDNA to satisfy meiotic HR requirements.

## Results

### In MEIOB-deficient mice, DMC1 loss correlates with BLM accumulation

Consistent with previous reports, RAD51 and DMC1 foci numbers decreased prematurely in *Meiob*-deficient mice (*SI*, S1*A,B*) (19, 30). Similar results have been described in *Spata22*-deficient rodents (30) To better understand this defect, we measured the colocalization of DMC1 and RAD51 (Fig. 1*A* and *SI*, S1*A, B*). We found that DMC1 and RAD51 foci showed equivalent levels of colocalization on *Meiob^+/+^* and *Meiob^-/-^* meiotic chromosomes (~70% of RAD51 foci colocalized with DMC1 foci on chromosome axes during prophase I; Fig. 1*B* and *SI*, S1*A, B*). Combined with published data indicating that RAD51 and DMC1 foci numbers are not altered by the absence of MEIOB during early steps of meiotic HR, our results suggest that both recombinases are properly loaded onto DSB sites in the absence of MEIOB. Costaining for DMC1 and RPA2 revealed that while RPA foci numbers decreased after the disappearance of DMC1 in *Meiob^+/+^* spermatocytes, RPA foci numbers remained high in *Meiob^-/-^* spermatocytes (Fig. 1*C*). These observations confirm that meiotic HR does not progress properly in the absence of MEIOB.

**Figure 1.**
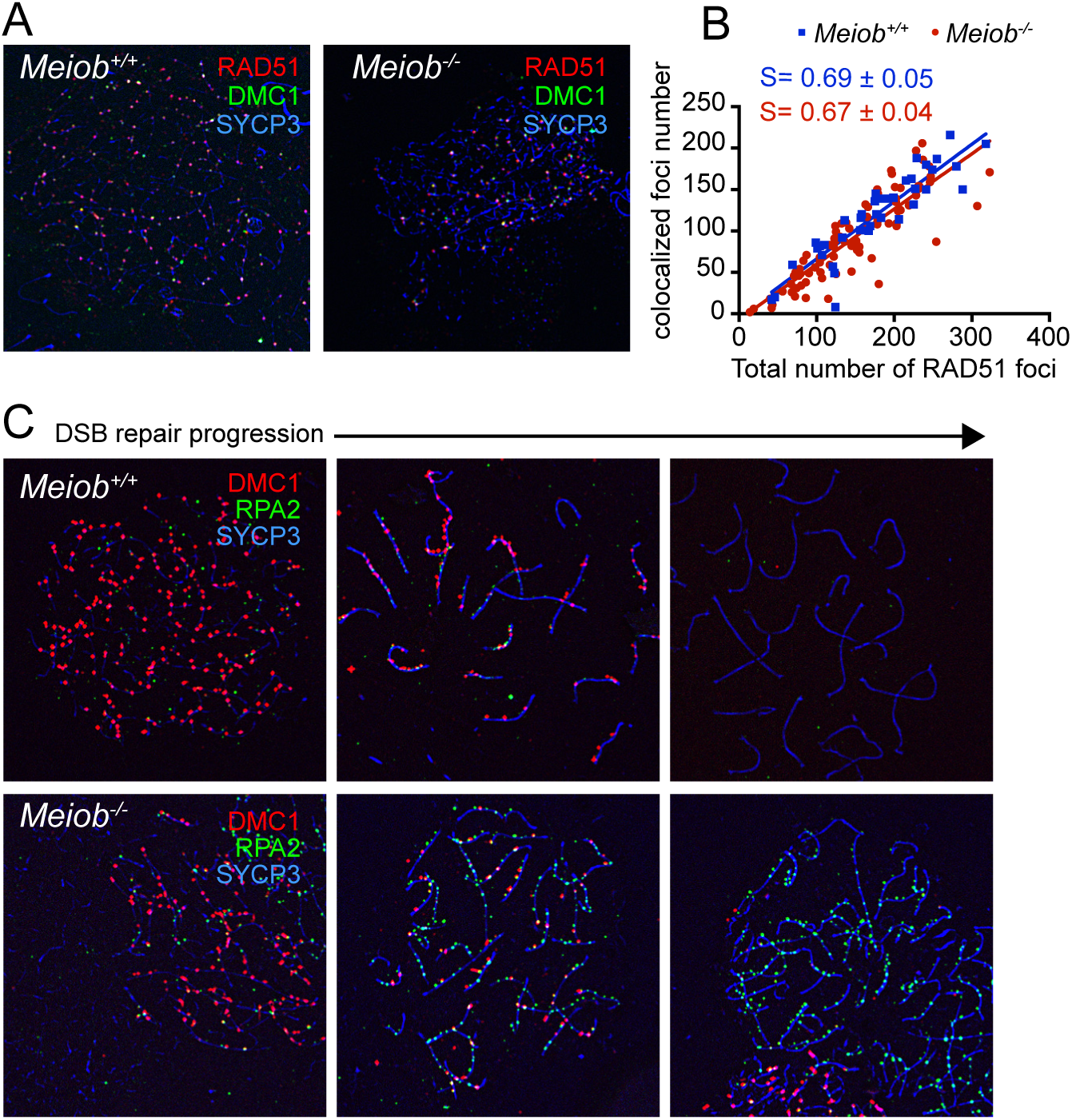
Unrepaired DSBs accumulate after the loss of RAD51 and DMC1 recombinases in MEIOB-deficient spermatocytes. (A) Representative wild-type and *Meiob^-/-^* spermatocyte nuclei immunostained for RAD51 (red) and DMC1 (green), and for the lateral axis marker SYCP3 (blue). (B) Plot representing the coexistence of RAD51 and DMC1 foci as a function of the number of RAD51 foci per meiotic cell in *Meiob^+/+^* or *Meiob^-/-^. Meiob^+/+^*, n=42 cells from 2 mice; *Meiob^-/-^*, n=90 cells from 2 mice. Linear regression curves are shown for *Meiob^+/+^* (R^2^=0.82) and *Meiob^-/-^* (R^2^=0.76), S=slope of the linear regression with 95% intervals. (C) Representative wild-type and *Meiob^-/-^* spermatocyte nuclei immunostained for DMC1 (red), RPA2 (green) and SYCP3 (blue). The arrow highlights DSB repair progression during meiotic prophase. Accumulation of RPA2 staining during late stages of prophase in MEIOB-deficient spermatocytes is indicative of defective DSB repair.

To better characterize the premature removal of recombinases that occurs in the absence of MEIOB, we investigated Bloom syndrome RecQ-like helicase (BLM) localization in *Meiob^-/-^* spermatocytes. BLM and its orthologs have been shown to play a role in the regulation of HR intermediates (6, 31–33). *In vitro* assays have demonstrated that human BLM can displace RAD51 from ssDNA, and BLM-deficient cells have been shown to accumulate RAD51 (34, 35). On *Meiob^+/+^* meiotic chromosomes, most BLM foci are faint and are located outside of chromosome axes; only a few bright foci are located on the axes, and they are observed mostly during zygotene and early pachytene (Fig. 2*A*, upper panel). In contrast, in the absence of MEIOB, bright BLM foci accumulate on chromosome axes during zygotene-like and pachytene-like stages (Fig. 2*A*, lower panel). Interestingly, BLM foci colocalized with RPA2 on chromosome axes, indicating that BLM accumulates at unrepaired DSBs in the absence of MEIOB (Fig. 2*A*). To determine whether the formation of BLM foci correlates with the disappearance of recombinases, we costained and quantified DMC1 and BLM foci (Fig. 2*B* and *C*). Their distribution differs in *Meiob^+/+^* and *Meiob^-/-^* spermatocytes. In spermatocytes where DMC1 foci numbers are similar, BLM foci numbers are on average higher in *Meiob^-/-^* compared to WT. Moreover, high BLM foci numbers coincide with decreased DMC1 foci numbers in *Meiob^-/-^* spermatocytes, whereas WT spermatocytes display a low number of BLM foci irrespective of DMC1 foci number. Interestingly, we also observed BLM accumulation in *Spata22*-mutant spermatocytes (*SI* Fig. S2). These results suggest that in the absence of MEIOB or SPATA22, BLM accumulation may trigger DMC1 removal.

**Figure 2.**
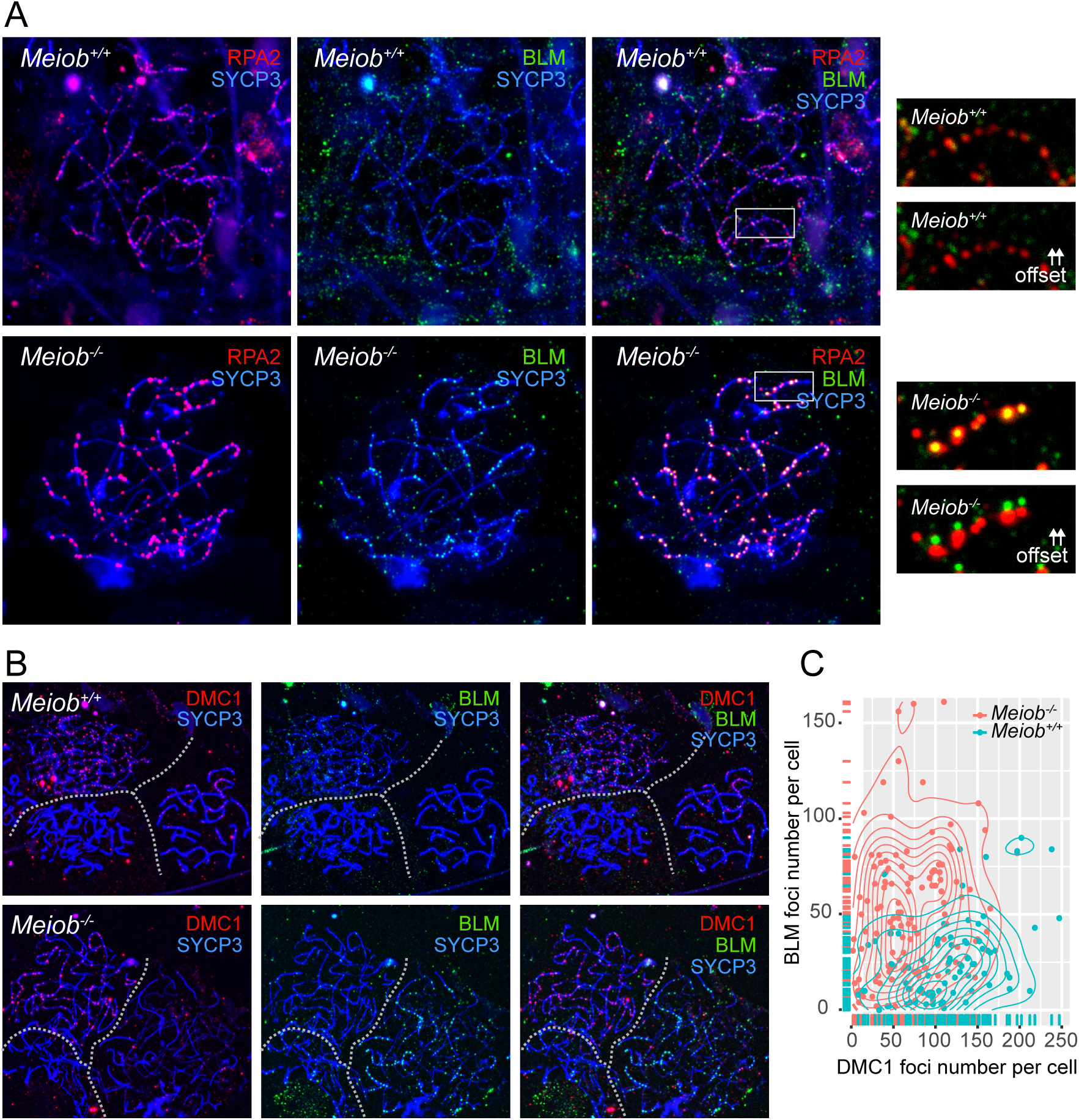
BLM accumulates on meiotic chromosomes in the absence of MEIOB. (A) BLM foci colocalize with RPA in the absence of MEIOB. Representative chromosome spreads of *Meiob^+/+^* and *Meiob^-/-^* spermatocyte nuclei immunostained for RPA2 (red), BLM (green) and SYCP3 (blue). Enlarged views of regions highlighted by white boxes are shown on the right, with the red and green channels merged (up) or offset (down). (B) Chromosome spreads of *Meiob^+/+^* and *Meiob^-/-^* spermatocytes immunostained for DMC1 (red), BLM (green) and SYCP3 (blue). *Meiob^+/+^* cells show little BLM staining on meiotic chromosome axes, whereas *Meiob^-/-^* spermatocytes often exhibit BLM accumulation. *Meiob^-/-^* cells that present BLM accumulation show faint DMC1 staining. White dashed lines separate individual cells. (C) Scatter plot density with marginal distribution of BLM and DMC1 foci numbers on the Y- and X-axes, respectively. Each point represents one meiotic cell. *Meiob^+/+^* (n=81 cells from 4 mice); *Meiob^-/-^* (n=120 cells from 4 mice).

### Aberrant “off axis” recombinase staining occurs in the absence of MEIOB

While studying DMC1 and RAD51 dynamics, we observed aberrant recombinase staining in *Meiob^-/-^* mice that had been recently crossed with an outbred strain. In WT, DMC1 and RAD51 form discrete round foci on chromosome axes (Fig. 3*A* left panel and *SI*, Fig. S1*A*). In *Meiob^-/-^* however, in addition to foci that appear normal, we observed aberrant “off axis” DMC1 staining with an elongated rather than round shape (Fig. 3*A* right panel). The quantity of “off axis” staining per cell and the number of cells with at least one “off axis” staining are highly variable between mice, but accumulate as meiotic prophase progresses; as many as 45 “off axis” staining structures are observed per cell during the pachytene-like stage, and they are present in more than 50% of spermatocytes in some *Meiob^-/-^* mice (Fig. 3*B*). Similar “off axis” DMC1 staining is also observed in *Spata22*-mutant mice with mixed genetic background (*SI* Fig. S2). Although we sometimes observed elongated DMC1 staining along the chromosome axis in WT mice, we failed to detect “off axis” elongated staining. Moreover, among the numerous spreads we analyzed, we did not observe “off axis” elongated staining with RAD51. The elongated DMC1 “off axis” staining is always connected to the axis and is often associated with a RAD51 focus on the axis (>90%). Interestingly, analysis by structure-illumination microscopy (SIM) revealed that the “off axis” elongated DMC1 structures are comprised of juxtaposed DMC1 foci that are sometimes interrupted or accompanied by a RAD51 focus (Fig. 3*C* and 3*D*). Together these results suggest that MEIOB and SPATA22 may restrain extensive DMC1 loading.

**Figure 3.**
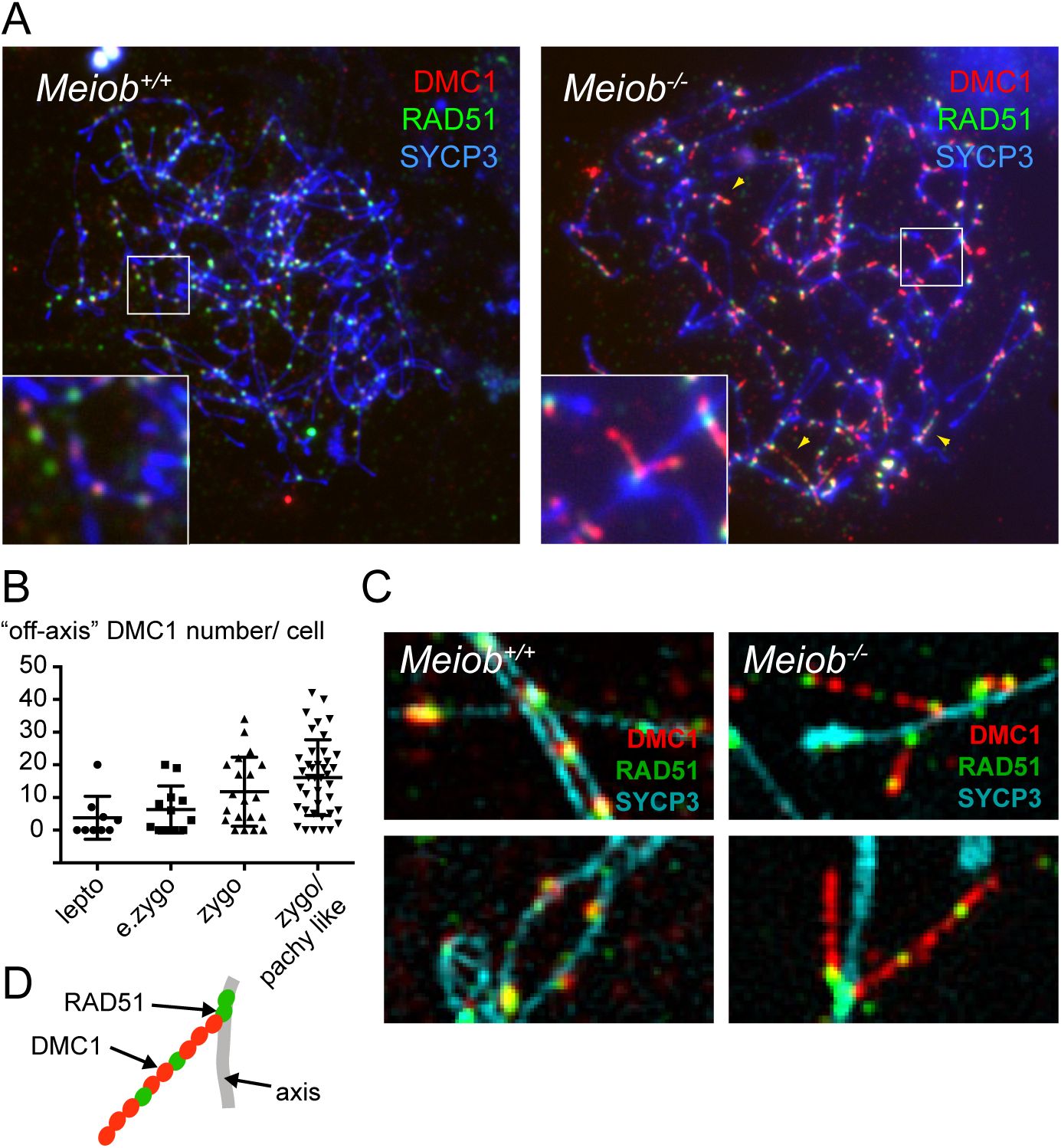
DMC1 shows aberrant “off axis” staining in the absence of MEIOB. (A) Representative chromosome spreads of spermatocytes from *Meiob^+/+^* or *Meiob*^-/-^ immunostained for DMC1 (red), RAD51 (green) and SYCP3 (blue) observed by epifluorescence microscopy. Yellow arrowheads highlight “off-axis” DMC1 staining observed in *Meiob^-/-^*. Representative regions containing RAD51 and DMC1 foci (white boxes) are enlarged (B) Quantification of “off-axis” DMC1 staining during different meiotic prophase stages in *Meiob*^-/-^ (n=2 mice). Each point represents one meiotic cell. One elongated DMC1 staining starting form the axis and pointing outside the axis was counted as one “off-axis” staining. Lines represent mean ± SD. (C) Representative SIM images showing DMC1 (red) and RAD51 (green) staining in chromosome spreads of spermatocytes from *Meiob^+/+^* or *Meiob^-/-^* mice. SYCP3 (cyan) stains the lateral axis. “Off-axis” DMC1 staining in the absence of MEIOB is often comprised of juxtaposed DMC1 foci interrupted by a RAD51 focus. (D) Schematic representation of elongated “off-axis” DMC1 staining in the absence of MEIOB. These structures are frequently associated with a RAD51 focus on the axis.

### MEIOB-SPATA22 directly binds and condenses RPA-coated ssDNA

Previous studies have demonstrated that MEIOB interacts with RPA subunits and ssDNA (19, 26, 29). We hypothesized that MEIOB could influence RPA-coated ssDNA properties to modulate HR dynamics during meiosis. RPA is a heterotrimer. Therefore, the intensity of any one subunit, as detected by immunostaining, should be proportional to the intensity of the other subunits. To test this, we compared the intensity of RPA1 and RPA2 foci formed on DNA lesions induced by camptothecin exposure in human cultured cells (Fig. 4*A*). As expected, RPA1 and RPA2 appeared as bright, colocalized foci with correlated intensities (Fig. 4*A*, offset and graph). Interestingly, during wild-type meiosis, RPA1 foci intensity did not correlate with RPA2 foci intensity, whereas during *Meiob*^-/-^ meiosis, good correlation was observed with a similar profile to that in cultured cells (Fig. 4*B*). These results indicate that the presence of MEIOB influences RPA detection.

**Figure 4.**
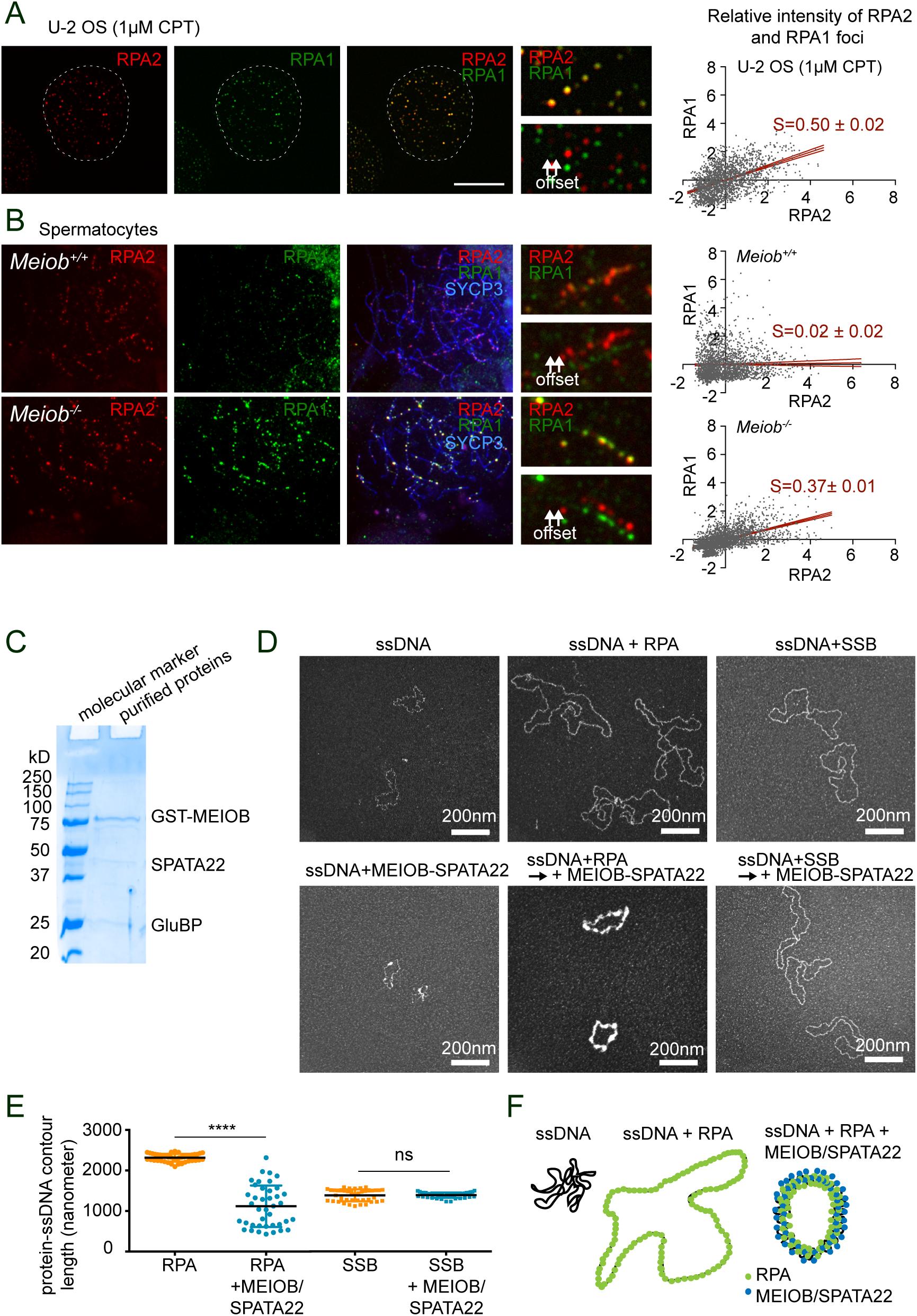
MEIOB-SPATA22 condenses RPA-coated ssDNA. (A) Representative U-2 OS cell treated with 1 μM camptothecin and immunostained for RPA1 (green) and RPA2 (red). The white dashed line highlights the nucleus. Right panels show magnified views of representative regions containing RPA1 and RPA2 foci (merges (up) and offsets (down)). The relative intensity of RPA1 foci as a function of the relative intensity of RPA2 foci is presented in the plot after Z-score standardization. Each point represents one focus (n=2023 foci, S=slope of the linear regression with 95% intervals shown in red, R^2^=0.24). (B) Representative wild-type and *Meiob^-/-^* spermatocyte nuclei immunostained for RPA1 (green) and RPA2 (red). Right panels show magnified views of representative RPA1 and RPA2 foci-containing regions (merges (up) and offsets (down)). RPA intensities are plotted as described for panel A. *Meiob^+/+^* (two mice, n=2131 foci, R^2^=0.0004); *Meiob^-/-^* (two mice, n=2763 foci, R^2^=0.29). (C) Purified GST-MEIOB-SPATA22 analyzed by SDS-PAGE (1.2μg of protein). GluBP: endogenous glutathione-binding protein from insect host cells that coelutes with MEIOB-SPATA22. (D) Transmission electron microscopy (TEM) images of circular ssDNA (ΦX174 virion) incubated alone, with RPA, or with SSB are shown in the top panel. Bottom panel shows images of ssDNA incubated with MEIOB-SPATA22, with or without prior incubation with RPA/SSB (approximately one RPA per 20 nucleotides, one SSB tetramer per 10 nucleotides, and one MEIOB-SPATA22 per 400 nucleotides). The incubation of MEIOB-SPATA22 with ssDNA forms heterogeneous bright structures, suggesting that MEIOB-SPATA22 forms aggregates in the presence of ssDNA. RPA covers the circular ssDNA to form a homogeneous complex. The addition of MEIOB-SPATA22 to preformed RPA-ssDNA complexes condenses the RPA-coated ssDNA. (E) Quantification of protein-coated ssDNA contour length in the presence of RPA or SSB alone, or with the addition of MEIOB-SPATA22. Lines depict mean ± SD, n=42 molecules for RPA, n=40 molecules for RPA+MEIOB-SPATA22, n=41 molecules for SSB and n=42 molecules for SSB+MEIOB-SPATA22; **** represents p< 0.0001 (two tailed, Mann-Whitney test); ns: nonsignificant. (F) Model showing ssDNA covered with RPA or with RPA and MEIOB-SPATA22. RPA binds ssDNA and prevents the formation of secondary structures. MEIOB-SPATA22 condenses RPA-coated ssDNA.

To gain insight into the role of MEIOB in modulating RPA behavior, we attempted to purify MEIOB and its essential partner SPATA22 for *in vitro* assays. Despite numerous attempts, we failed to purify MEIOB or SPATA22 individually, but we achieved copurification by GST-pull down after their coexpression in insect cells (Fig. 4*C*). We compared protein-coated ssDNA containing RPA and/or MEIOB-SPATA22 by transmission electron microscopy (TEM). Under saturating conditions, human RPA uniformly coats circular ssDNA (Fig. 4*D*). In contrast, MEIOB-SPATA22 formed aggregates when added to circular ssDNA (Fig. 4*D*). When MEIOB-SPATA22 was added to preformed RPA-coated ssDNA, the circular nucleoprotein complex became thicker and its contour length decreased (Fig. 4*D* and *E*). No aggregates were observed under these conditions. Importantly, the addition of MEIOB-SPATA22 did not modify preformed nucleoprotein complexes of ssDNA and SSB, a bacterial ssDNA-binding protein (Fig. 4*D* and *E* for quantification). These results suggest that MEIOB-SPATA22 interacts specifically with RPA-coated ssDNA to change its conformation (Fig. 4*F*).

To verify that MEIOB-SPATA22 and RPA subunits coexist on ssDNA, we performed electrophoretic mobility shift assays and immunoblot analyses. Increasing amounts of MEIOB-SPATA22 were incubated with a Cy5-labeled 46-mer ssDNA that either was or was not preincubated with a saturating amount of RPA. The altered migration of ssDNA induced by the addition of proteins was visualized on non-denaturing acrylamide gels (Fig. 5*A*). The addition of increasing amounts of MEIOB-SPATA22 in the absence of RPA induced multiple band shifts with the persistence of free ssDNA (Fig. 5*B*, short and long migration, lane 3 to 5). This result shows that the MEIOB-SPATA22 complex binds ssDNA and thus retains biochemical activity. However, due to experimental limitations we were unable to obtain a concentration of MEIOB-SPATA22 that would allow us to reach saturating conditions in this assay (i.e. free ssDNA remains after addition of the highest amount of MEIOB-SPATA22, Fig. 5*B*, short migration, lane 5). Interestingly, when increasing amounts of MEIOB-SPATA22 were added to ssDNA preincubated with RPA, we detected an accumulation of molecules with lower mobility compared to RPA-coated ssDNA molecules (asterisk in Fig. 5*B*, long migration, lanes 6 to 9). To verify that MEIOB and SPATA22 are present along with RPA in these super-shifted bands, the migration products were transferred to a membrane and immunostained for MEIOB, SPATA22 and RPA subunits. Because the antibodies directed against MEIOB and SPATA22 were raised in the same species, samples were loaded twice for independent immunostaining of MEIOB and SPATA22. Immunostaining for MEIOB or SPATA22 alone without ssDNA revealed that the two proteins migrate as a complex (Fig. 5*C*, lane 1). They do not migrate as a single discrete band but show a long smear, suggesting that the complex may adopt different organizational structures (Fig. 5*C*, lane 1). MEIOB and SPATA22 are both detected on shifted ssDNA in the absence of RPA, which shows that MEIOB and SPATA22 bind ssDNA (Fig. 5*C*, lanes 3 to 5). In the presence of RPA, MEIOB and SPATA22 are both detected on molecules with the lowest mobility (asterisk in Fig. 5*C*, lanes 7 to 9). Immunostaining of RPA subunits RPA1, RPA2 and RPA3 shows that they are also present on these molecules, supporting the idea that MEIOB-SPATA22 and RPA coexist on ssDNA (Asterisk on Fig. 5*C*, lane 7 to 9). As expected, RPA subunits are detected on shifted ssDNA observed in the absence of MEIOB-SPATA22 (Fig. 5*C*, lane 6, shifts 1 and 2). The fact that these molecules persist after the addition of increasing amounts of MEIOB-SPATA22 suggests that a large portion of the RPA-coated ssDNA remained unbound by MEIOB-SPATA22 under our experimental conditions (Fig. 5*C*, lanes 7 to 9, shifts 1 and 2, *SI*, Fig. S3).

**Figure 5.**
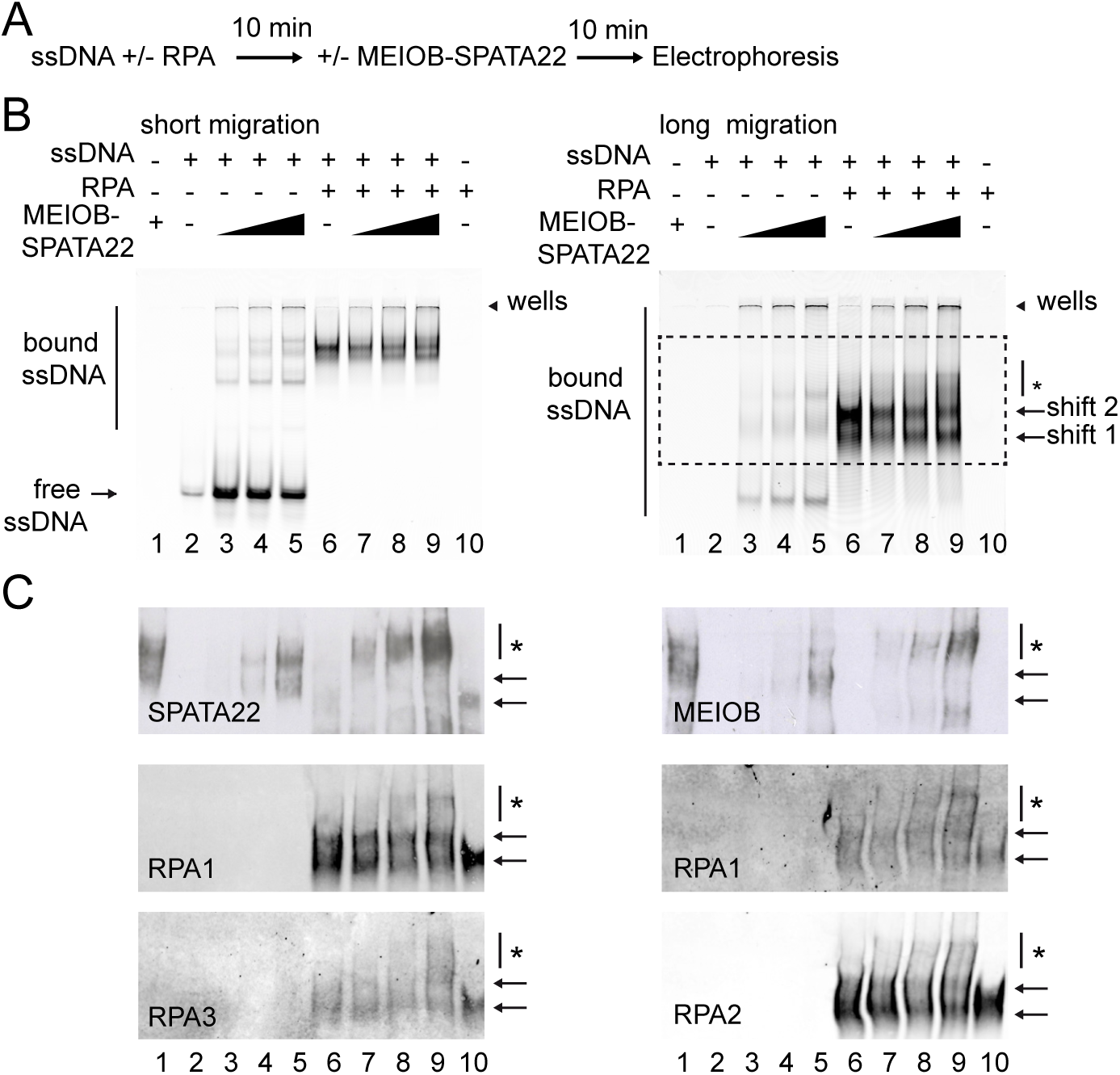
MEIOB-SPATA22 coexists with RPA on ssDNA. (A) Schematic representation of the gel shift assay performed. A Cy5-labeled 46-mer ssDNA was incubated with or without RPA. Increasing amounts of MEIOB-SPATA22 was added to the reactions, and protein-ssDNA complexes were resolved by nondenaturing polyacrylamide gel electrophoresis. (B) Gel shift assay with a short gel migration time (left panel) and a longer migration time (right panel) for the same gel. The addition of RPA to ssDNA induces two shifted bands (arrows 1 and 2). The addition of MEIOB-SPATA22 to preformed RPA-coated ssDNA induces additional smeared shifted bands (* in right panel). (C) Proteins within the dotted rectangle in panel B (long migration) were detected using anti-RPA1, anti-RPA2, anti-MEIOB, or anti-SPATA22 antibodies. Full-size scans are presented in *SI Appendix*, Fig. S3. RPA1, in the left and right panels, was detected with two different antibodies due to experimental limitations. Data are representative of two independent experiments. MEIOB and SPATA22 comigrate with ssDNA and RPA subunits in the smeared shifted bands (*).

### MEIOB and SPATA22 share structural features with subunits of the RPA complex

Crystal structures have been solved for portions of the RPA complex from various species (36–39). We previously showed that MEIOB is a paralog of RPA1 (19, 40), the largest subunit of the trimeric RPA complex. Indeed, MEIOB possesses three OB-fold containing domains (OBCDs) consisting of an OB-fold (five-or six-stranded closed β-barrels) and supplementary secondary structures. We modeled the structure of human MEIOB using the protein structure prediction server RaptorX (41), which selected RPA1 as the best template (Fig. 6*A*). Similarities between MEIOB and RPA1 were revealed by superimposition of RPA1 OBCDs 1 to 3 from *Ustilago maydis* (UmRPA1) (named DBDA, DBDB, and DBDC in (39)) and the predicted structure of human MEIOB (HsMEIOB) (Fig. 6*A*). Structure prediction revealed the presence of a zinc ribbon and a helix-turn-helix motif in the predicted OBCD3 of MEIOB. These specific features are also present in the OBCD3 of RPA1 (Fig. 6*A*). Given the strong structural similarities between MEIOB and RPA1, we asked whether SPATA22 also possesses similarities to RPA subunits. The RaptorX server selected HsRPA2 as the best template to model the structure of SPATA22 and predicted the presence of a putative OBCD in the C-terminal part of human SPATA22 (Fig. 6*B*, left panel)(41). Superimposition of the predicted HsSPATA22 structure and the HsRPA2 crystal structure highlights the structural similarities between SPATA22 and RPA2 (Figure 6*B*, right panel) (42).

**Figure 6.**
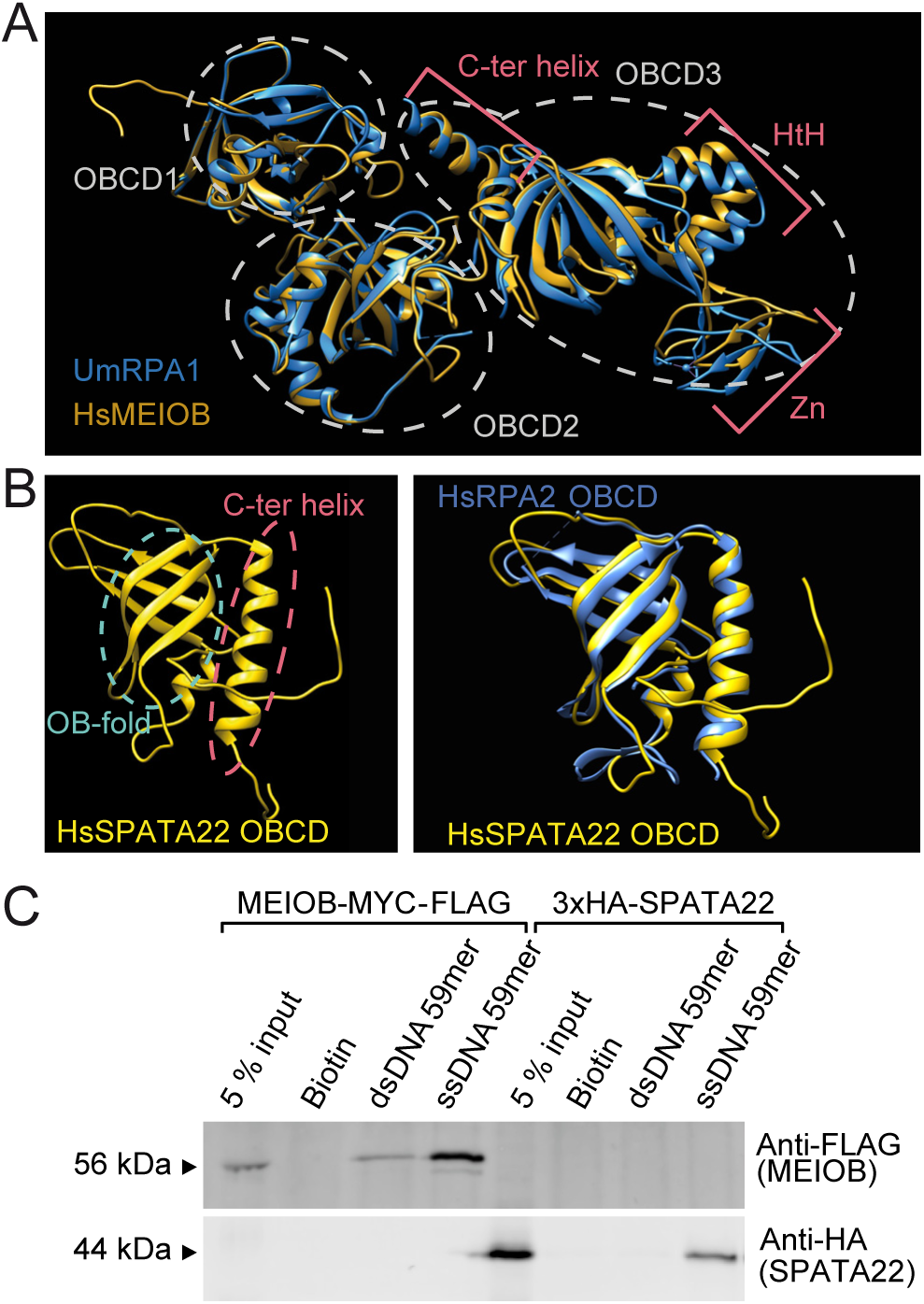
MEIOB and SPATA22 have OB-fold-containing domains and structural similarities to RPA. (A) Superimposition of RPA1 OB-fold-containing domains (OBCDs) 1 to 3 from *Ustilago maydis* (UmRPA1) (39) and the predicted structure of human MEIOB (HsMEIOB). The predicted structure of HsMEIOB was obtained by alignment to UmRPA1 OBCDs 1 to 3, which is the most complete RPA1 OBCDs 1 to 3 structure available (39). Gray dotted circles highlight the OBCDs of UmRPA1 and HsMEIOB. Red brackets highlight the C-terminal helix (C-ter helix), the helix-turn-helix (HtH), and the zinc ribbon domains (Zn) in OBCD3 of MEIOB and RPA1. (B) Left panel: ribbon representation of the predicted OBCD of human SPATA22 (HsSPATA22). The cyan dotted circle represents the OB-fold, and the red dotted circle highlights the -helix flanking this OB fold. Right panel: superimposition of ribbon representations of the OBCDs of HsRPA2 and HsSPATA22 (predicted). (C) Tagged versions of full-length human MEIOB or SPATA22 were expressed in a cell-free system and mixed with biotin alone or with biotinylated double-stranded or single-stranded 59-mer DNA (59-mer dsDNA or 59-mer ssDNA, respectively) and precipitated with streptavidin-coupled magnetic beads. Precipitated proteins were detected using anti-FLAG (MEIOB) or anti-HA (SPATA22) antibodies. The data are representative of three independent experiments.

Given the similarities between SPATA22 and RPA subunits, we next asked whether SPATA22 interacts with ssDNA. Human versions of HA-tagged-SPATA22 or MYC-FLAG-tagged-MEIOB were expressed *in vitro* in a cell-free system and pulled down with either a biotinylated 59-mer ssDNA or a biotinylated 59-mer dsDNA (Fig. 6*C*). MEIOB was preferentially pulled down by ssDNA, as previously reported (19, 26), as was SPATA22 (Fig. 6*C*). This observation suggests that both MEIOB and SPATA22 bind ssDNA.

### MEIOB and SPATA22 interaction resembles the interactions between RPA subunits

To study MEIOB and SPATA22 interactions, we took advantage of the similarities that these proteins share with RPA subunits. Previous structural studies revealed that the three RPA subunits interact via OBCDs that form the minimum component of their interaction or trimerization core (38, 39, 43–46). Given that similar domains are present in MEIOB and SPATA22, we hypothesized that these may be involved in MEIOB-SPATA22 interactions.

To test this hypothesis, we constructed wild-type and truncated forms of tagged human MEIOB and tagged human SPATA22 and transiently expressed them in cultured HEK293 cells (Fig. 7*A*). The interactions between these two proteins were tested by coimmunoprecipitation (co-IP) using antibodies directed against their tags, with strong nuclease treatment to avoid aberrant interactions due to oligonucleotide bridges (see Materials and Methods). MEIOB and SPATA22 immunoprecipitated with each other, consistent with previous data obtained with murine proteins (Fig. 7*B*, lane 6 and *C*, lane 6; see *SI*, Fig. S4*A* and *B* for input). This demonstrates that the MEIOB-SPATA22 interaction is conserved in humans (26, 29). We next examined truncated versions of MEIOB and SPATA22 that lack C-terminal OB-fold containing domains (Fig. 7*A*). Co-IPs showed that C-terminal OBCDs of MEIOB (MEIOB OBCD3) and SPATA22 (SPATA22 OBCD) are required to ensure their interaction (Fig. 7*B*, lane 7, 7 *C*, lane 7, and *SI*, Fig. S4*A*, and *B*). These results are consistent with recent work by Xu et al., who expressed murine constructs of MEIOB and SPATA22 containing a series of deletions to identify the regions required for their interaction (29). Next, we questioned whether the OBCDs of MEIOB and SPATA22 are sufficient to ensure their interaction, as suggested by their structural similarity to RPA. By coexpressing and immunoprecipitating (i) OBCD3 of MEIOB with full-length SPATA22, (ii) OBCD of SPATA22 with full-length MEIOB, or (iii) OBCD3 of MEIOB and OBCD of SPATA22, we observed that OBCD3 of MEIOB and OBCD of SPATA22 are sufficient to ensure the interaction between MEIOB and SPATA22 (Fig. 7*B*, lane 8, 7 *C* lane 8, 7*D* lane 6). A published analysis of the crystal structure of RPA highlighted the role of OBCD C-terminal helixes in RPA complex formation (38). Interestingly, the predicted structures of MEIOB and SPATA22 suggest that these proteins contain similar helixes flanking their C-terminal OB folds (Fig. 6*A* and *B*). Deletion of the SPATA22 C-terminal helix abolishes MEIOB- SPATA22 interaction (Fig. 7*C*, lane 9 and *SI*, Fig. S4*B*). However, deletion of the MEIOB C-terminal helix did not abolish this interaction (Fig. 7*B*, lane 9 and *SI*, Fig. S4*A*). These results are expected, as truncating part of the C-terminal domain of RPA1, which contains the helix, is insufficient to abolish its interaction with RPA2 (43). Altogether, our observations demonstrate that similar to RPA, the OBCD3 of MEIOB and the OBCD of SPATA22 are crucial for their interaction (Fig. 7*E*).

**Figure 7.**
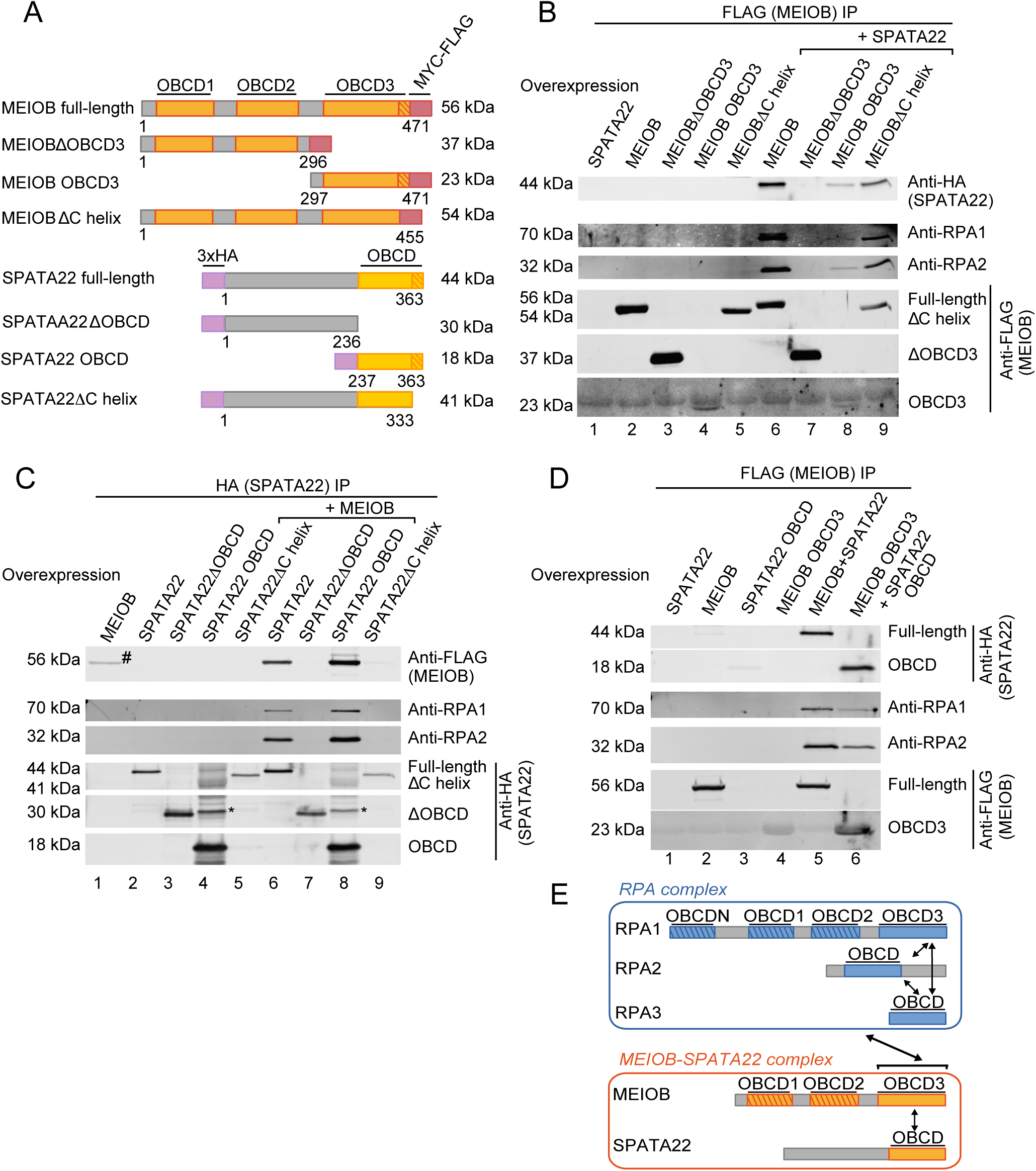
Interactions between MEIOB and SPATA22 share similarities with interactions between RPA subunits, and MEIOB-SPATA22-interacting domains ensure its interaction with RPA. (A) Schematic representation of full-length and truncated forms of human tagged MEIOB and human tagged SPATA22 used in this study. OBCDs of MEIOB are in orange, OBCD of SPATA22 is in yellow, C-terminal helixes are shown as striped boxes, and MYC-FLAG and 3xHA tags are shown in red and purple, respectively. (B) MEIOB and/or SPATA22 were transfected into HEK293 cells and MEIOB was immunoprecipitated (FLAG (MEIOB) IP) using anti-FLAG antibody. Full-length and truncated forms of MEIOB were expressed alone (lanes 2 to 5) or coexpressed with SPATA22 (lanes 6 to 9). SPATA22, RPA1 and RPA2 immunoprecipitated proteins were detected using anti-HA, anti-RPA1 and anti-RPA2 antibodies, respectively. Tagged full-length MEIOB, ΔOBCD3, OBCD3 and ΔC helix were detected with anti-FLAG and displayed expected sizes of 56, 37, 23 and 54 kDa. The data are representative of three independent experiments. (C) MEIOB and/or SPATA22 were transfected into HEK293 cells and SPATA22 was immunoprecipitated (HA (SPATA22) IP) using anti-HA antibody. Full-length or truncated forms of SPATA22 were expressed alone (lanes 2 to 5) or coexpressed with MEIOB (lanes 6 to 9). MEIOB, RPA1 and RPA2 immunoprecipitated proteins were detected using anti-FLAG, anti-RPA1 and anti-RPA2 antibodies, respectively. Tagged full-length SPATA22, ΔOBCD, OBCD and ΔC helix were detected with anti-HA and displayed expected sizes of 44, 30, 18 and 41 kDa. (*) represents a nonspecific band or a posttranslational modification of SPATA22 OBCD that displays a molecular weight greater than 30 kDa. (#) MEIOB is retained slightly in a nonspecific manner on the anti-HA column. The data are representative of minimum two independent experiments. (D) MEIOB OBCD3 and SPATA22 OBCD cooperate to interact with RPA. MEIOB and/or SPATA22 were transfected into HEK293 cells, and MEIOB was immunoprecipitated (FLAG (MEIOB) IP) using anti-FLAG antibody. Full-length SPATA22 and full-length MEIOB expressed alone (lanes 1 and 2) or coexpressed (lane 5). SPATA22 OBCD and MEIOB OBCD3 expressed alone (lanes 3 and 4) or coexpressed (lane 6). MEIOB, SPATA22, RPA1 and RPA2 immunoprecipitated proteins were detected using anti-FLAG, anti-HA, anti-RPA1 and anti-RPA2 antibodies, respectively. The data are representative of two independent experiments. (E) Model of interactions between MEIOB, SPATA22 and RPA. MEIOB and SPATA22 interactions recapitulate interactions between RPA subunits and involve their C-terminal OBCDs. Furthermore, MEIOB and SPATA22 collaborate to interact with RPA. Striped boxes represent OBCDs that are not involved in complex formation.

### MEIOB and SPATA22 cooperate to interact with endogenous RPA

MEIOB and SPATA22 have been shown to interact with RPA subunits (26, 29). Moreover, MEIOB and SPATA22 colocalize with RPA1 and RPA2 on chromosome spreads of spermatocytes (26, 30) (19). To test whether MEIOB and SPATA22 interact with the RPA complex, we studied their interactions with endogenous RPA in human cells, where the three RPA subunits are expected to exist as a tripartite complex. When expressed alone, MEIOB and SPATA22 did not co-IP with endogenous human RPA1 or RPA2 (Fig. 7*B*, lane 2 and *C*, lane 2 and *SI*, Fig. S4*A* and *B* for input). However, when they were coexpressed, MEIOB and SPATA22 coimmunoprecipitated with RPA1 and RPA2, irrespective of whether MEIOB or SPATA22 was immunoprecipitated (Fig. 7*B*, lane 6 and *D*, lane 6 and Fig. S4*A* and *B*). We were not successful in detecting RPA3 in co-IP experiments due to technical limitations. We further demonstrated that OBCD of SPATA22 and OBCD3 of MEIOB are necessary (Fig. 7*B*, lane 7, and Fig. 7*C*, lane 7) and sufficient (Fig. 7*D*, lane 6) for MEIOB and SPATA22 to interact with each other (see above) and to interact with endogenous RPA1 and RPA2 (Fig. 7*E* and *SI*, Fig. S4 for input) (29).

### Discussion

MEIOB was recently identified as a meiosis-specific factor that is essential for meiotic prophase. Here we used cellular and molecular approaches to characterize the role of MEIOB and its essential partner SPATA22. Our findings suggest that MEIOB and SPATA22 work together with RPA to ensure proper meiotic recombination.

We and others previously showed that DMC1 and RAD51 recombinases are loaded but prematurely dissociate from meiotic chromosomes in the absence of MEIOB or SPATA22 (19, 30). In the absence of MEIOB, RAD51 and DMC1 colocalization is maintained, reinforcing the idea that they are normally loaded onto chromosomes. However, numerous RPA foci persist after DMC1 dissociation from chromosomes, indicating that recombination does not progress properly. The BLM helicase accumulates in cells with low DMC1 foci number, suggesting that BLM may be responsible for the clearance of recombinases from arrested meiotic recombination intermediates. Moreover, BLM colocalizes with RPA, indicating that recombination intermediates are arrested at a step that contains ssDNA. We hypothesize that impaired HR in the absence of MEIOB triggers BLM accumulation at ssDNA, thus inducing the eventual removal of the recombinases and RPA accumulation.

Interestingly, we find that RPA subunit detection is influenced by the presence of MEIOB; RPA1 and RPA2 foci intensity correlate in cultured cells and *Meiob^-/-^* spermatocytes, but do not correlate in wild-type spermatocytes. A tempting explanation is that MEIOB and RPA interaction influences the accessibility of antibodies directed against RPA subunits.

Our biochemical approaches with purified proteins show that MEIOB-SPATA22 and RPA coexist on ssDNA and form highly compacted protein-coated ssDNA. While the band shift assays demonstrate their coexistence on ssDNA, we cannot rule out the possibility that there are mixed individual complexes containing MEIOB, SPATA22, and/or RPA2 or RPA3. The existence of such complexes can also explain the disparity in RPA1 and RPA2 immunodetection in *Meiob^+/+^* spermatocytes. Our data suggest that MEIOB-SPATA22 interacts with RPA to confer specific properties to ssDNA-containing recombination intermediates.

We previously proposed that MEIOB plays a role in modulating presynaptic recombinogenic filaments, as MEIOB was loaded onto meiotic chromosomes in *Dmc1^-/-^* mice that are unable to perform strand invasion of the homolog (19). Several RPA-interacting proteins have been shown to modulate the ssDNA-binding activity of RPA (47). For example, BRCA2, with the help of DSS1, locally displaces RPA and overcomes RPA inhibition of RAD51 and DMC1 loading onto ssDNA (48–51). One possibility is that MEIOB-SPATA22 limits RPA displacement by reducing its accessibility to specific factors and prevents extensive polymerization of recombinases. Recent biochemical data demonstrated that DMC1 displaces more RPA from ssDNA compared to RAD51, and that they form homotypic extended filaments (52). These data support the idea that MEIOB-SPATA22 may function to reinforce RPA-ssDNA binding to prevent excessive DMC1 loading during meiosis. These findings could explain why aberrant “off axis” extended DMC1 staining is sometimes observed in absence of MEIOB. The fact that this phenomenon is only observed in mice with heterogeneous genetic backgrounds suggests that DNA polymorphisms between parental chromosomes may play a role, and could explain why this staining pattern has not been previously described in studies performed with more inbred mice. Further experiments will be needed to understand the involvement of polymorphism. Aberrant loading of RAD51 and DMC1 has been proposed to occur sometime on dsDNA (53) (23, 54). The fact that we did not observe aberrant “off axis” staining for RPA suggests that the DMC1 “off axis” staining may involve dsDNA. Moreover, because most DMC1 “off axis” staining are accompanied by a RAD51 focus on the chromosome axes, this “off axis” staining is likely initiated at a DSB site and may elongate along chromosome loops. Previous studies analyzed either RAD51 or DMC1 alone, while we investigated both recombinases simultaneously by SIM microscopy. We therefore cannot exclude the possibility that the aberrant staining we observed is a general consequence of the accumulation of aberrant recombination intermediates.

In addition to its role in recombinogenic filament formation, RPA binds the displaced strand (D-loop) to facilitate strand invasion (12, 14). MEIOB, SPATA22 and RPA colocalize and form bright foci in late zygotene and early pachytene, which is consistent with their loading onto D-loops during meiotic recombination (19, 26). It is tempting to speculate that the compaction of RPA-coated ssDNA by MEIOB-SPATA22 provides D-loops with specific properties that are essential for meiotic recombination. These properties could influence D-loop stabilization, migration or extension. They could also change the affinity of RPA-coated ssDNA for partners. A tempting hypothesis would be that stabilizing the D-loop could favor second end capture and formation of double holiday junctions to ensure crossovers. The fact that MEIOB, SPATA22, and RPA foci persist longer than the recombinase foci is consistent with a potential role during maturation of D-loops.

We took advantage of the RPA structure to show that MEIOB and SPATA22 share structural features with RPA subunits. Moreover, we experimentally demonstrated that MEIOB and SPATA22 interact in an RPA-like manner. In particular, we showed that the last OBCD of MEIOB and the OBCD of SPATA22 are necessary and sufficient for their interaction; they constitute a minimum dimerization core that resembles the trimerization core of RPA. Recently, Xu et al. identified mutations in MEIOB and SPATA22 that abolished their interaction. Consistent with our results, these mutations fall within the OBCD3 of MEIOB and the OBCD of SPATA22, and likely perturb their folding (29).

We demonstrated that MEIOB and SPATA22 cooperate to interact with RPA and that their minimum dimerization core is sufficient for this interaction. Recently published data showed that murine MEIOB and SPATA22 cooperate to interact with murine RPA3 (29). Taken together, the MEIOB-SPATA22-RPA interaction may be mediated via an interaction between the MEIOB-SPATA22 dimerization core and the RPA3 subunit. Interestingly, when we expressed MEIOB or SPATA22 alone, we did not observe interactions with RPA1 or RPA2. These results differ from those of Xu et al., who identified interactions between MEIOB or SPATA22 expressed individually and RPA1 or RPA2 expressed individually, via regions outside of the dimerization core (29). These variations may be due to experimental differences. Xu et al. identified interactions between murine proteins overexpressed in human cells and tested RPA subunits individually, whereas we expressed human MEIOB and SPATA22 in the presence of the endogenous human RPA complex. Notably, Xu et al. did not coimmunoprecipitate endogenous human RPA, most likely due to the interactions being species-specific. Based on the study by Xu et al. and our work, we propose that MEIOB-SPATA22-RPA complex formation requires MEIOB and SPATA22 to interact, and the dimerization core formed (composed of C-terminal OBCDs) interacts with RPA via RPA3. Their interaction may be further stabilized by secondary independent contacts between RPA1 and/or RPA2 and regions of MEIOB and SPATA22 that are located outside of their dimerization core.

The interactions between MEIOB-SPATA22 and RPA suggest that these proteins may collaborate to form specialized, meiosis-specific protein-coated ssDNA, and MEIOB-SPATA22 likely provide specific properties required to accomplish meiotic recombination.

Materials and Methods

### ssDNA and dsDNA pulldown assay

ssDNA and dsDNA pulldowns were performed as described (19) with modifications; details are in *SI Materials and Methods*. Briefly, human SPATA22 and human MEIOB were expressed using TnT T7 Quick Coupled Transcription/Translation System (Promega). Binding reactions were performed with biotinylated ssDNA or dsDNA and Dynabeads M-280 Streptavidin (Life Technologies) in binding buffer (25 mM Tris-HCl, pH 7.5, 100 mM NaCl, 1 mM EDTA, 5 mg/ml BSA, 0.05% Tween 20, 10% glycerol, 1 mM ß-mercaptoethanol, complete protease inhibitor without EDTA (Roche)), washed in rinsing buffer (25 mM Tris-HCl, pH 7.5, 150 mM NaCl, 1 mM EDTA, 0.05% Tween 20, 10% glycerol, 1 mM ß-mercaptoethanol, complete protease inhibitor without EDTA (Roche)), and eluted in Laemmli buffer.

### Co-Immunoprecipitation

Co-IPs are detailed in *SI Materials and Methods*. HEK-293 cells were lysed in supplemented cell lysis buffer (Cell Signaling), and proteins bound overnight in binding buffer (20 mM Tris-HCl, pH 7.5, 150 mM NaCl, 10% glycerol, 1 mM EDTA, 1 mM beta-mercaptoethanol (except for HA IP), 10 mg/ml BSA, complete protease inhibitor without EDTA (Roche), 10 μM MG-132, 90 U/ml Benzonase nuclease (Novagen)). Additional Benzonase digestion was done with the following reagents: 90 U/ml Benzonase nuclease, 20 mM Tris-HCl, pH 8.0, 20 mM NaCl, 10% glycerol, 2 mM MgCl_2_, 1 mM beta-mercaptoethanol (except for HA IP), 10 mg/ml BSA and complete protease inhibitor without EDTA (Roche).

### Immunofluorescence on U2-OS cells

To remove free proteins and retain the chromatin-associated fraction, cells were permeabilized before fixation as described (55). Details are in *SI Materials and Methods.*

### Immunofluorescence on preparations of chromosome spreads

Chromosomes spreads are detailed in *SI Materials and Methods.*

## Acknowledgements

We thank members of the Laboratory of the Development of the Gonads for discussions and technical support, and are especially grateful to M.J Guerquin for data analysis. We thank members of the animal housing facility at iRCM, in particular V. Neuville and C. Joubert. We thank M.S. Wold for RPA plasmids and P. Drevet for development of the baculovirus system. We are especially grateful to M. Grelon, V. Borde and J. Haber for careful reading and constructive comments on the manuscript. This work was supported by ANR, INSERM, Fondation ARC, and La Ligue contre le Cancer. J.R. was supported by a CEA fellowship and La Ligue contre le Cancer. D.J. was supported by a Human Frontier Science Program fellowship.

